# Somatosensory signals from the controllers of an extra robotic finger support motor learning

**DOI:** 10.1101/2021.05.18.444661

**Authors:** E. Amoruso, L. Dowdall, M.T. Kollamkulam, O. Ukaegbu, P. Kieliba, T. Ng, H. Dempsey-Jones, D. Clode, T.R. Makin

## Abstract

Considerable resources are being invested to provide bidirectional control of substitutive and augmentative motor interfaces through artificial somatosensory feedback. Here, we investigated whether intrinsic somatosensory information, from body part(s) proportionally controlling an augmentation device, can be utilised to infer the device’s state and position, to better support motor control and learning. In a placebo-controlled design, we used local anaesthetic to attenuate somatosensory inputs to the big toes while participants learned to operate a toe-controlled robotic extra finger (Third Thumb) using pressure sensors. Motor learning outcomes were compared against a control group who received sham anaesthetic. The availability of somatosensory cues about the amount of exerted pressure generally facilitated acquisition, retention and transfer of motor skills, and performance under cognitive load. Motor performance was not impaired by anaesthesia when tasks involved close collaboration with the biological fingers, indicating that the brain could ‘close the gap’ of the missing pressure signals by alternative means, including feedback from other body parts involved in the motor task. Together, our findings demonstrate that there are intrinsic natural avenues to provide surrogate position information to support motor control of an artificial body part, beyond artificial extrinsic signalling.

## INTRODUCTION

The last three decades have seen dramatic advances in the development of interfaces that can extend, substitute or restore human upper-limb motor function. Such interfaces allow users (for example, amputees and tetraplegic patients) to operate assistive or augmentative robotic limbs by decoding intended movements at different levels along the motor system. Current technologies allow the extraction of motor commands from the user’s muscle activations and residual movements (e.g., myoelectric or body-powered prostheses) or directly from neuronal signals (Brain Machine Interfaces – BMI) (Bensmaia & Miller, 2014).

However, the functionality of these motor interfaces is restricted by the scarce sensory feedback available to the user, whose movements are typically guided mainly through visual monitoring. In natural conditions, motor control heavily relies on (often subconscious) somatosensory signals for tracking limb state and interactions with objects. Tactile signals from the skin mechanoreceptors are especially important for object manipulation, as they convey information about contact timing, size, and location, as well as the optimal amounts of pressure to exert (Johansson & Flanagan, 2009). Proprioceptive signals allow us to plan and guide the dynamics of limbs movements by informing us on current joint position and motion (Proske & Gandevia, 2012). The importance of somatosensory feedback for motor control is well evidenced by the debilitating motor deficits that characterize a total loss of touch and proprioception, as is the case in some acute sensory neuropathies (Gordon et al., 1995). Moreover, increasing evidence shows that the somatosensory system plays a fundamental role not only in the online movement planning and correction processes, but also in motor learning, i.e. the acquisition and consolidation of novel motor skills (Vidoni et al., 2010; see Ostry & Gribble, 2016, for a review).

In light of this, considerable resources are being invested to provide bidirectional sensorimotor control of prosthetic limbs (Bensmaia et al., 2020). Most of the progress in this work has involved providing information about contact between the device and objects (i.e., touch), either by providing tactile stimulation on a displaced skin surface (Antfolk et al., 2013; see also Hussain et al., 2015 for applications in augmentation), or by directly activating the neural pathways originally supporting the sensory function, e.g. the peripheral nerves (Dhillon and Horch, 2005; Raspopovic et al., 2014; Tan et al., 2014), the spinal cord (Chandrasekaran et al., 2020), or the somatosensory cortex (Romo et al., 1998; Tabot et al., 2013; see Bensmaia & Miller, 2014, for a review). Although the development of such tactile afferent interfaces is at a much earlier stage than their efferent counterparts and critical challenges remain (e.g., the quality of the evoked percepts), the benefits of artificial touch while using bionic limbs are being documented in both lab and home settings (Ortiz-Catalan et al., 2020; Graczyk et al., 2018). Conversely, very little progress has been made in reproducing proprioception to provide kinaesthetic feedback, which would allow the user to feel the prosthesis’ current configuration and state position without the cognitively taxing effort of constant visual monitoring. This omission is mainly due to difficulties in developing reliable systems for artificially providing the multifaceted proprioceptive signals, which in normal conditions are derived from a combination of multiple afferent channels (i.e., muscles spindle fibers, Golgi tendons, joint angle and cutaneous receptors), and to an incomplete understanding of the representation of this modality at the cortical level (Tomlinson & Miller, 2016). Owing to such technical and scientific considerations, attempts to restore (London et al., 2008; Suminsky et al., 2010; Dadarlat et al., 2015) or substitute (Blank et al., 2008; Schiefer et al., 2018; D’anna et al., 2019) proprioceptive feedback in artificial limbs haven’t so far been able to show consistent benefits for motor control.

The challenges currently posed by reproducing artificial forms of somatosensory, particularly proprioceptive, feedback prompt us to consider simplified approaches that could potentially take advantage of the intrinsic bodily signals emerging from the prosthesis control. For example, in the case of traditional myoelectric or body-powered prostheses, when motor commands are sent out by the bodily controller (residual arm and shoulder, respectively), users receive a range of somatosensory inputs. These are not only sensations on the patch of skin interfacing the prosthetic actuator that can inform about contact with objects, but could also include, theoretically, indirect cues about the prosthesis state and position. Surrogate position cues could be extracted from the body part which is proportionally controlling the device, through muscle activations or active movements and pressure exertion. The benefits of such physiologically built-in feedback systems, which may circumvent the need for complex substitutionary or restorative interfaces, have long been highlighted in clinical and rehabilitative settings (Hirsch & Klasson, 1974; Simpson, 1974), yet mostly unexplored. For example, it has been speculated that such intrinsic, although limited, forms of feedback may contribute to the choice of body-powered prostheses over more sophisticated ones (e.g. myoelectric), due to the increased sensory information that is received on the user’s body when operating the device (e.g. from extending the cable/harness with the shoulder) (Antfolk et al., 2013). To our knowledge, however, whether this naturally available sensory information is actually being harnessed by the motor system for supporting the control of artificial limbs has not been thoroughly investigated or quantified.

To this end, supernumerary (extra) limbs provide an innovative experimental model for isolating the role of intrinsic forms of somatosensory feedback on artificial limb motor control. The Third Thumb (Dani Clode Design, Figure 1A-B) is a 2 degrees of freedom (DOF) robotic sixth digit which is worn on the hand (receiver) and wirelessly operated by the big toes (controllers). The Thumb is proportionally controlled using pressure sensors strapped underneath the big toes. To operate the device, the user needs to vary the pressure applied on the sensors using each of the big toes (right for flexion/extension and left for abduction/adduction). The somatosensory inputs received by the toes during device control (through pressure exertion) could be potentially leveraged to infer the Thumb’s state and position (device proprioception) to better support motor control and learning (Wolpert & Ghahramani, 1995).

**Figure 1.**
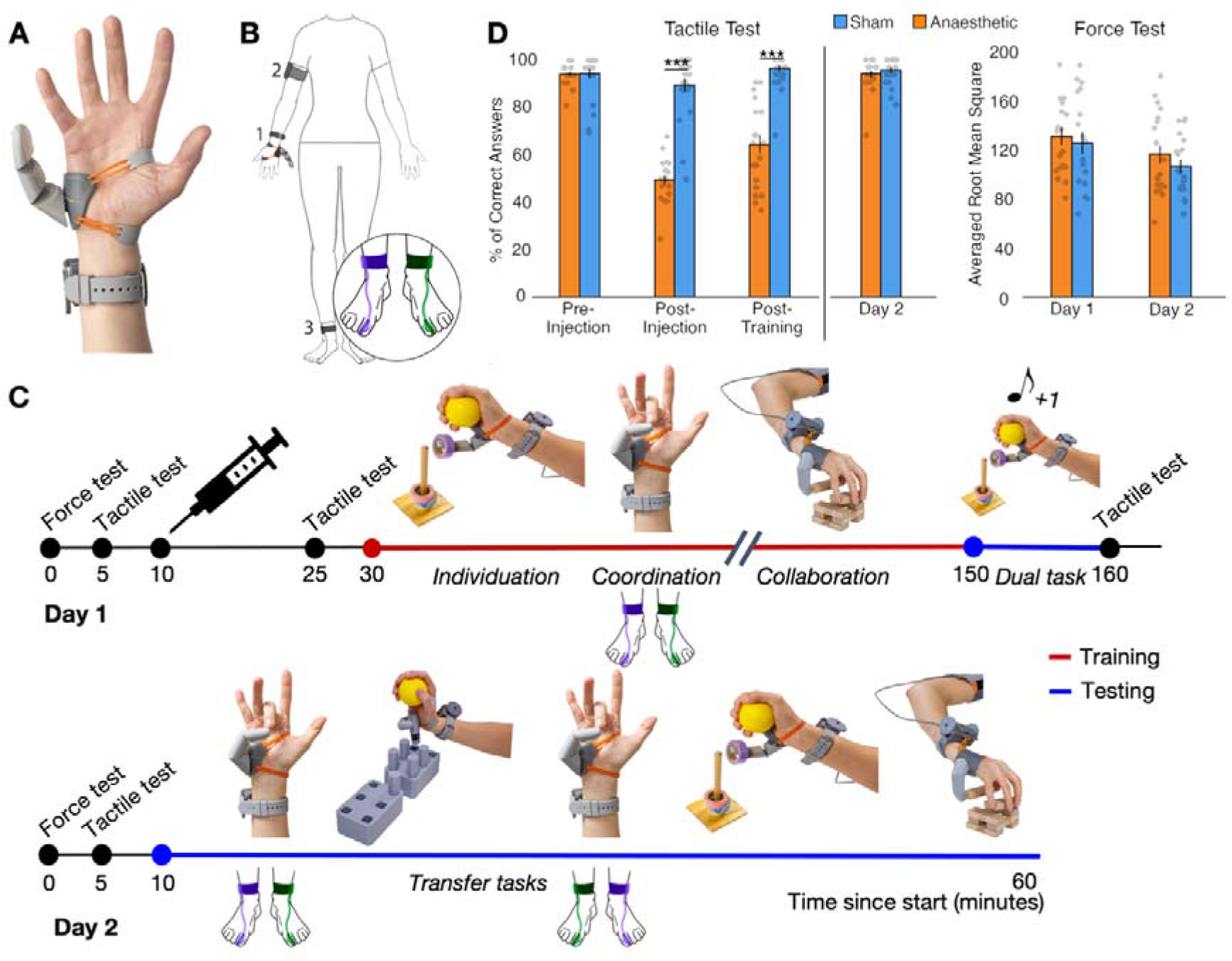
Experimental design. **A-B)** The Third Thumb is a 3D-printed robotic extra digit. Mounted on the side of the palm (1), the Thumb is actuated by two servomotors (fixed to a wrist band), allowing for independent control over flexion/abduction. The Thumb is powered by (2) an external battery, strapped around the arm and wirelessly controlled by (3) two pressure sensors strapped underneath the participant’s big toes. The colours purple and green represent the sensors controlling the flexion/extension and abduction/adduction movements, respectively. **C)** Timeline depicting the course of the study over the two days. The syringe on Day 1 represents the anaesthetic/sham intervention. Training with the Third Thumb is represented in red and testing in blue. Tasks used for training and testing are displayed on the timeline: Individuation – of the Third Thumb (picking up, transporting and repositioning tape rolls) while keeping the biological hand occupied; Coordination – between the biological fingers and Third Thumb: opposing the tips of the fingers to the tip of the Thumb; Collaboration – between the biological hand and Thumb: lifting and transporting wooden bars to build a tower; Dual task – increasing the cognitive load on the individuation task, by simultaneously conducting a primary working-memory numerical task; Transfer tasks – varying task demands relative to training (1) picking up, transporting and repositioning pegs for Third Thumb individuation and (2) the coordination task with the flexion/extension and abduction/adduction pressure sensors swapped. The training tasks were completed in a randomised order while the testing tasks were completed in the fixed order shown in the timeline. **D)** Participants across groups showed similar performance in the baseline tactile acuity test and confirmed expected changes in tactile sensitivity following the injection intervention according to group assignment. Participants also showed similar performance across groups at baseline and on Day 2 on the Force test, used to assess baseline motor ability and determine the pseudo-randomised group allocations. The bars depict group means; error bars represent standard error of the mean. Individual dots correspond to individual subjects’ scores. *Asterisks denote significant effect of Timepoint at ***p<0.001.*

Healthy individuals were first trained to use the Third Thumb and then tested on a series of motor learning outcomes over the course of two days (Figure 1C). To modulate intrinsic somatosensory contributions to the Thumb’s state and position, we used anaesthetic injections to attenuate pressure feedback to the big toes in a test group prior to training. This procedure prevents most tactile signals from the toes skin/Golgi tendon receptors, while leaving motor function (enabled by foot/ankle muscles) and proprioception (arising from the ankle’s muscle spindles) mostly unaffected. Training performance and learning outcomes were compared against a control group, who received sham injections. We compared participants’ ability to acquire, retain and transfer (Kantak & Winstein, 2012) their motor skills to infer the potential benefits of the controllers’ pressure feedback on motor learning. We hypothesised that, by receiving more complete somatosensory feedback about the amount of pressure exerted with the toes, the Sham group would be able to develop a more accurate internal model of the Third Thumb state and position, allowing more effective motor learning. We also hypothesised that reducing intrinsic position cues will incur more attention to monitor Thumb performance through complementary sensory cues (e.g. visual). As a consequence, the Anaesthetised group would show more deficits in device control when a further cognitive load is added to motor tasks (Poldrack et al., 2005).

## METHODS

### Participants

50 right-handed participants were recruited for the study. Exclusion criteria included allergies to local anaesthetics, needle phobia and history of neurological or psychiatric illness. 4 participants dropped out of the study due to a vasovagal response to the injections and 2 participants did not proceed beyond the injections due to an unsuccessful numbing effect (see: Anaesthetic Intervention). The final participant group included 44 participants with 22 participants in the ‘Anaesthetised’ group (11 female, mean age = 22.91, SD= 3.85, range = 18 to 35 years) and 22 participants in the ‘Sham’ group (10 female, mean age = 23.45, SD = 3.60, range = 18 to 33 years). Participants were pseudo-randomly allocated to either the ‘Anaesthetised’ or ‘Sham’ group (see: Sampling Validation Tests). Due to updates to the study design after initial data collection, only 40 participants (20 in each group) completed the Dual Task and only 17 participants in the Sham group and 15 participants in the Anaesthetised group completed the retention test for the Collaboration and Individuation tasks. Due to technical issues during data acquisition, pressure sensors recordings from 2 participants in the Anaesthetised group and 5 in the Sham group could not be analysed. All participants provided their informed consent prior to participation. Ethical approval for the study was granted by the UCL Ethics committee (Project ID: 12921/001).

### Third Thumb

The Third Thumb (Dani Clode Design, London, UK) is a robotic extra finger that attaches to the ulnar side of the right hand (see Figure 1A) and is wirelessly operated through pressure sensors strapped underneath the big toes. The Thumb has two degrees of freedom that allow a corresponding proportional control: applying pressure to left big toe sensor causes an adduction/abduction movement, whilst pressure on the right one causes a flexion/extension movement (Figure 1B). The pressure applied by each big toe when the Thumb was switched on during the experiment was recorded and used for off-line analyses (see section: Toes Pressure Analysis). Both pressure sensors contained an SD-card that logged the position of the Thumb’s servomotors (range 10-175 degrees for flexion/extension, range 10-80 degrees for abduction/adduction). The position of the motor was linearly proportional to the pressure applied by the participant. This relationship was fixed for all participants in such a way that the same value of pressure was always associated with the same servo position. When using the Third Thumb, participants placed their big toes on a small platform of a footrest, to minimise somatosensory feedback from the other toes or neighbouring foot surface while controlling the Thumb.

### Anaesthetic intervention

All participants received the same deafferentation protocol (Dempsey-Jones et al., 2019). The intervention varied only in the substance injected, which depended on group assignment. Each anaesthetic intervention required an injection of 2.5ml of 2% Lidocaine Hydrochloride and 2.5ml of 0.5% Bupivacaine Hydrochloride. Lidocaine Hydrochloride is a fast-acting anaesthetic ensuring an almost immediate numbing effect, while Bupivacaine Hydrochloride is a long-lasting anaesthetic ensuring the numbness lasted for the length of the Day 1 sessions. The Sham injection consisted of 3ml of 0.9% Sodium Chloride. The injections were administered on both big toes by medically trained professionals. A 25–gauge sterile needle was inserted into the base of the dorsolateral aspect of the big toe bilaterally and the solution was injected as the needle was withdrawn, achieving a sensory deafferentation of the entire toe. This procedure prevents afferent sensory inputs from the toe, whereas motor function (mostly enabled by foot/ankle muscles) is largely preserved. In the absence of an acceptable numbing effect, indicated by a score of ~50% in the tactile acuity check (see section: Sampling Validation Tests), the medical professional made a clinical decision on whether to administer further injections. To create a sham effect, participants in both groups were informed that they were receiving local anaesthetic, but that the effects were variable, and they therefore may not subjectively perceive a complete anaesthetic effect.

### Experimental Design

The study was conducted over the course of 2 days, using the same design across both groups. An overview of the experimental time course is shown in Figure 1C. Motor and tactile validation tests on the big toes (black line in Figure 1C) were performed at several different timepoints (see Figure 1D for outcomes). Prior to the pharmacological intervention, force and tactile tests were performed to establish baseline performance, followed by the anaesthetic/sham injections. A post-injection tactile acuity test was performed by one experimenter. A second experimenter, in charge of administering the subsequent tests, was blinded to the group assignment. A further tactile acuity test was conducted at the end of Day 1. The same force and tactile acuity tests were administered at the beginning of Day 2.

Third Thumb training (red line in Figure 1C) was conducted on Day 1 over 3 consecutive sessions, each comprising of 3 different training tasks (Individuation, Coordination, Collaboration; see section: Training Tasks), with task order randomised across participants. Tasks were performed over 10-minute blocks.

Testing tasks (blue line in Figure 1C) were performed at the end of Day 1 (Dual Task) and on Day 2 (retention and transfer tests). Task order was fixed on Day 2. Participants completed one 10-minute block of the Coordination task, followed by one block of an adapted individuation task (Individuation Transfer), and one block of an adapted coordination task (Coordination Transfer). Finally, single blocks of the Collaboration and Individuation tasks were completed.

### Sampling Validation Tests

#### Tactile acuity

Orientation discrimination tests (adapted from Tong et al., 2013) were conducted at four different timepoints (before the injections, immediately after the injections, at the end of Day 1, at the beginning of Day 2; Figure 1D) to assess baseline acuity and the effectiveness of the anaesthetic intervention. A Touch Test^®^ two-point discriminator with a 13mm spacing was presented to the glabrous surface of the distal pad of each big toe for ~2-3 seconds in one of two randomly assigned orientations, with a total of 16 trials per test and inter-trial intervals of ~5 seconds. Participants verbally reported the perceived orientation using a two-alternative forced choice (‘down’ or ‘across’). The outcome measure was the percentage of correct answers.

This data was used to confirm similar group performance during baseline and that the anaesthetic intervention was successful at attenuating somatosensory inputs throughout Day 1. Due to violations of the normality assumption, as well as the presence of several tied values and the lack of true continuity in the data, group differences in tactile acuity were analysed using percentile bootstrap independent-samples tests (see: Statistical Analyses). Before receiving the injections, both groups showed equal tactile acuity on the big toes (mean accuracy Anaesthetised = 93% ± 4.6%, Sham = 94.0% ± 9.6%; bootstrapped 95% CI [−4.12%, 4.47%], p=.935). On Day 2, the Sham and Anaesthetised groups performed similarly and close to ceiling (mean accuracy Anaesthetised = 94% ± 6.6%, Sham = 95.6% ± 5.7%; bootstrapped 95% CI [−5.40%,1.82%]; p=.370), suggesting that the numbing effects had effectively washed out by the following day.

Next, we confirmed that the injections of local anaesthetic affected tactile acuity relative to the sham injections. Following the injections, tactile perception in the Anaesthetised group dropped to chance level (mean accuracy = 49.3% ± 8.8%), whereas performance of the Sham group was relatively high (mean accuracy = 89.2% ± 13.3%), resulting in a significant group difference immediately after the injections (bootstrapped 95% CI [−46.05%, −33.21%], p<.001). A similar effect was also recorded at the end of Day 1 (mean accuracy Anaesthetised = 64.3% ± 17%, Sham = 96.3% ± 5.9.6%; bootstrapped 95% CI [−39.43%, −24.44%], p<.001).

#### Force

A force test was administered at the very beginning of each day and repeated on each big toe in order to assess baseline motor abilities and determine the pseudo-random group allocation of the participants. Participants pressed down on a force sensor, taped to their big toe, to control the vertical direction of a horizontally-moving dot displayed on the laptop screen. Their aim was to maintain the appropriate force to hit a series of bars, whose height was determined using 20%, 50% and 80% of the maximum force the participant could apply. The force test consisted of four trials. Each trial contained 6 bars displayed simultaneously on the screen (each bar 20%, 50% or 80% of the participant’s maximum applied force). The bars of each trial were arranged in a way that either formed an up/down staircase (20% – 50% – 80% – 80% – 50% – 20%) or in a random order. Participants completed two identical up/down trials and two non-identical randomised trials. The outcome measure was the root mean square error from the ideal force required to maintain the moving dot at the top of each bar, averaged across all four trials. This data was used to confirm that baseline motor abilities were matched across groups and determine group allocation. The first 20 participants were randomly allocated to the Sham or Anaesthetised groups. The group means of the outcome measure were then calculated and participants thereafter assigned to groups based on their performance on the force test, to ensure that baseline motor abilities remained matched. Independent samples t-tests indicated that both Anaesthetised and Sham groups performed similarly in the force test at baseline on Day 1 (t(38)=0.36, p=.720) and on Day 2 (t(38)=0.22, p=.824).

### Training Tasks

Specific tasks were selected to probe the range of motor skills characterizing unimpaired Third Thumb motor control (Kieliba et al., 2021; Figure 1C). Retention of the trained tasks was additionally measured on Day 2.

#### Individuation

In the individuation task participants had to work on the fine motor control of the Third Thumb, while not relying on any support from the hand. Using only the Third Thumb, participants were required to pick up 6 tape rolls one-by-one and stack them on to a pole, while holding a foam ball to occupy their biological fingers (Figure 1C). Each block lasted for either 10 minutes or 10 trials, whichever came first. A trial lasted for however long it took the participant to stack all 6 rolls. If a given trial was still ongoing when 10 minutes had passed, the participant was allowed to complete it for up to an additional 5 minutes. The dependent variable was the time taken to stack all 6 tape rolls averaged across each block.

#### Coordination

To monitor the ability to coordinate the Thumb in synchrony with the hand, we used a finger opposition task (Meraz et al., 2018; Kieliba et al., 2021; Figure 1C). Participants were seated in front of a computer screen that displayed task stimuli. They were instructed to move the Thumb to touch the tip of a randomly specified finger of the augmented hand. A MATLAB script was used to randomly select a target finger (thumb, index, middle, ring or little) and to display the finger name on the computer screen. The experimenter manually advanced the program to the next stimulus when the participant successfully touched the tip of the target finger with the Third Thumb (hit), or when a wrong finger had been touched (miss). Participants were instructed to attempt to make as many successful hits as possible within a 1-minute trial, and completed a total of 10 trials per block. The outcome measure was the average number of hits completed in each block.

#### Collaboration

To measure continuous collaboration of the Third Thumb with the other fingers, participants were asked to build a 2×2 wooden bar tower with the augmented hand (Figure 1C). To do this they had to pick up 2 bars at a time from the table, using the Third Thumb in collaboration with a finger to hold or support one of the bars and two fingers to hold the other. Each block consisted of 10, 1-minute trials, with the dependent variable being the number of 2-bar floors built in 1 minute averaged across each block.

### Testing tasks

The testing tasks were variations of the training tasks, designed to examine the automaticity and flexibility of motor learning under different task requirements.

#### Dual task

In order to evaluate if pressure feedback from the Third Thumb controllers is associated with a reduced need for cognitive control over performance (i.e. with increased automaticity), participants were asked to complete a dual (motor and numerical) task at the end of Day 1. The task was adapted from previous studies, showing that numerical cognition impacts motor performance while controlling a virtual prosthetic arm (Witteveen et al., 2012) or a brain-computer interface (Guthrie et al., 2019), but does not impact Third Thumb control (Kieliba et al., 2021). The task involved completing an extra 10-minute block of the Individuation task, with a simultaneous counting task. Participants were instructed to complete the Individuation task as described above, but they now had to also complete a simultaneous counting out loud task, as their primary task. At the beginning of each trial a random number would be presented, then a series of high (550 Hz) and low (250 Hz) pitch tones would be sounded at random intervals (between 2 and 6 seconds) in a randomised order. Participants were instructed to add 1 to the current number after hearing a high tone, and subtract 1 from the current number after hearing a low tone. After each mathematical operation, participants were instructed to verbally respond with the resulting number. The outcome measure was the time taken to complete the secondary Individuation task, under the interference effects of the primary numerical task.

#### Coordination Transfer task

This task was aimed at probing transfer of learning by modulating task demands from training. The transfer version of the Coordination task was conducted using the same procedures as described above, but with the force sensors controlling the Third Thumb swapped onto the opposite feet. This resulted in having to adapt to a new mapping between the controlling movements of the toes and the Third Thumb responses, i.e. pressing with the left toe for a flexion/extension movement and with the right toe for adduction/abduction.

#### Individuation Transfer task

This task aimed to get participants to transfer the fine motor techniques developed in the Individuation task to a distinct setting. The task was inspired by grooved pegboard tasks widely used to assess motor functioning and dexterity in clinical neuropsychology (Rabin et al., 2005). A brick of 6 3-D printed pegs was placed to the participants’ left and a brick of 6 holes for the pegs to fit in was placed to their right. Participants were instructed to pick up each peg using only the Third Thumb and plant it in a designated hole, while holding a foam ball to occupy their biological fingers. They were instructed to move the pegs in a set order, starting from the right back peg, ending on the left front peg. The block was completed when all 6 pegs had been placed, or when 15 minutes had passed, whichever came first. Participants completed one block of this task. The outcome measure was the time taken to complete the block.

### Toes motor control analysis

As a follow-up analysis, we investigated potential differences in how the two groups used the big toe pressure sensors to control the Third Thumb at the very first stages of motor learning. We focused on performance during Block 1 of the Collaboration task. The data corresponding to the first block of the Collaboration task was identified using the timestamps and manual logs and was then imported into MATLAB. The average servomotor position (linearly dependent on the force applied) for both the flexion/extension and the abduction/adduction sensors for the duration of the Collaboration task was extracted for each participant. This was then normalised by dividing the average value by the maximum servomotor angle (respectively, 175 and 80 degrees). The total time spent making a flexion/extension or abduction/adduction movement was then calculated by first identifying the timestamps at which participants applied force, moving the servomotor out of its baseline position of 10 degrees. This was then used to find the proportion of task time corresponding to each of the movements. Using the same approach, we also calculated the time spent making bilateral movements (moving both servomotors at the same time), as a proportion of the total movement time.

### Statistical analyses

To identify violations of the normality assumption Shapiro-Wilk’s values were inspected. Within- and between-participants comparisons in task performance across acquisition, retention and transfer of coordination and collaboration skills were assessed using mixed analysis of variance (ANOVA). Significant interactions were followed up with confirmatory comparisons using independent and paired samples t-tests, where appropriate. For repeated-measurement factors with more than two factor levels, if the assumption of sphericity was violated, Huynh–Feldt corrected p-values are reported. Due to violations of normality, data from the individuation tasks, the tactile acuity tests and the toes pressure sensors were analysed using independent-samples percentile bootstrap tests, which do not require any assumption on the underlying distribution of the data. For each variable of interest, we calculated the difference in group means on each iteration (10000) and estimated a 95% confidence interval and p-value of the difference in group means from the resulting distribution. On all datasets, data points were removed separately for each task and/or block if they exceeded the outlier threshold of 2 standard deviations above or below the group mean. Less than 2% of data points across all tasks were removed for this reason. All parametric tests were performed on SPSS 25 (IBM, Chicago, Illinois), percentile bootstrap tests on RStudio 1.3 (RStudio PBC, Boston, Massachusetts).

## RESULTS

### Faster acquisition of robot individuation skills

We first considered the role of pressure feedback while controlling the Third Thumb individually, i.e. independently from the rest of the hand. Here, without any complementary information from the biological fingers about successful interaction with objects, task success is dependent on the development of dexterous motor control with the Third Thumb alone. We found that under these conditions, the anaesthetic intervention impacted the early stages of motor learning. Due to violations of the assumption of normality, we used bootstrapping approaches. To obtain a measure of the improvement on the task for both groups through Day 1 training, we subtracted the score (time taken to stack all the rolls) of Block 3 from Block 1, akin to the interaction term of a mixed ANOVA (Figure 2A). A bootstrapped independent-samples test performed on the obtained difference score was significant (bootstrapped 95% CI [28.297, 144.501], p=.003). Additional bootstrapped tests revealed a significant group difference in performance at Block 1 [Mean Anaesthetised (N=20) = 262s, Mean Sham (N=21) = 175s; bootstrapped 95% CI [13.216, 158.814], p=.018], but only marginally at Block 3 [Mean Anaesthetised (N=22) = 78s, Mean Sham (N=21) = 72s; bootstrapped 95% CI [−21.455, 29.160], p=.064], Therefore, the Anaesthetised group showed initial deficits on performance when first approaching this dexterous task, but eventually reached a similar skill level to the Sham group by the end of the training day.

**Figure 2.**
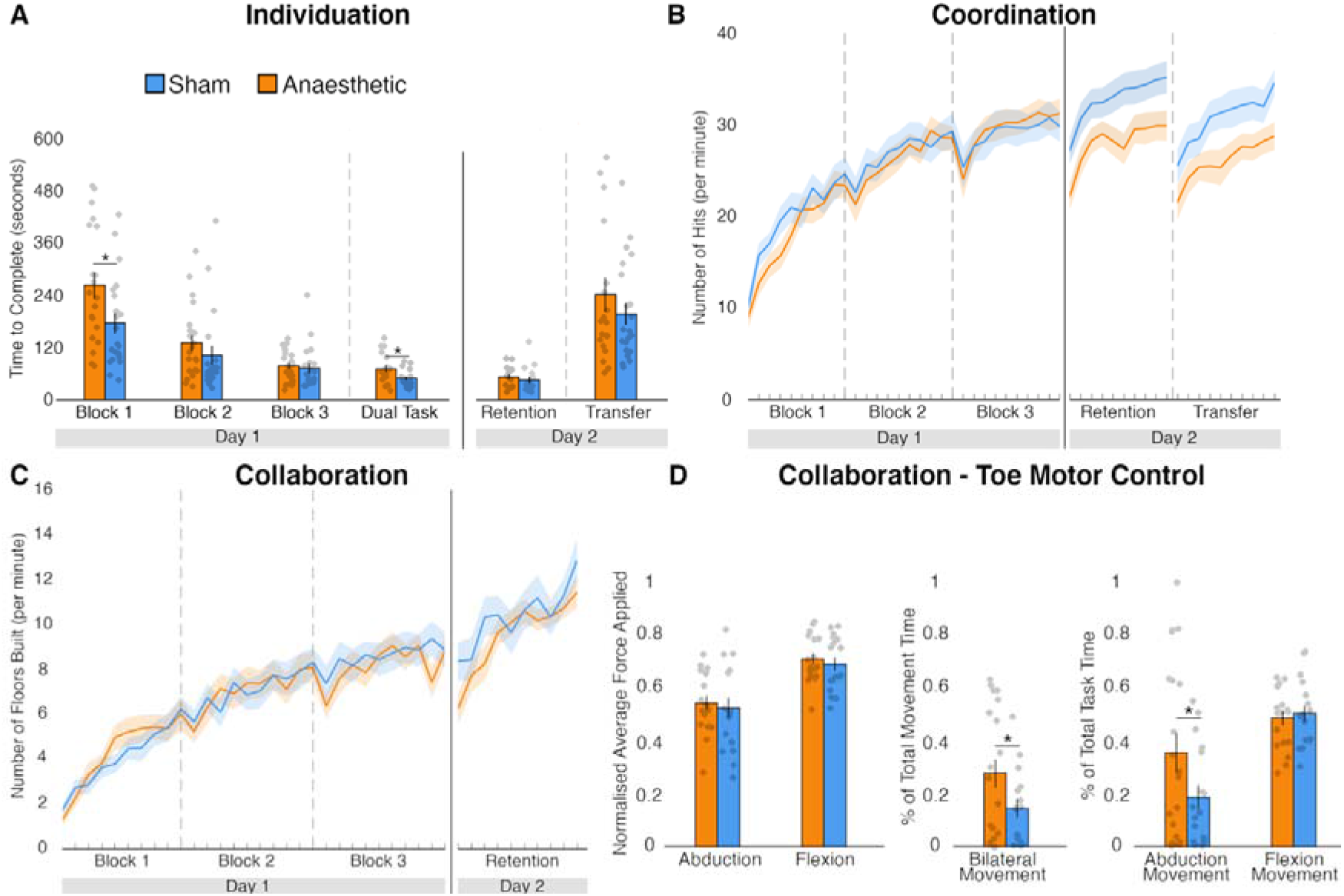
Task outcomes. **A)** In the Individuation task, the Anaesthetised group had significant deficits at the beginning of training and when a further cognitive load was added (Dual Task), but groups performed similarly in the retention and transfer tests of Day 2. **B)** In the Coordination task, despite similar Day 1 training performance, the Anaesthetised group showed significant deficits in the Day 2 retention and transfer tests. **C)** Both groups showed similar performance throughout the Collaboration task. **D)** Toe motor control analysis for Block 1 of the Collaboration task: no group differences in the amount of force applied; the Anaesthetised group used significantly more bilateral movements than the Sham group; the Anaesthetised group used significantly more adduction/abduction movements throughout the task, whereas there was no difference for flexion/extension movements. The bars depict group means; error bars represent standard error of the mean. Individual dots correspond to individual participants’ scores. *Asterisks denote significant effects at * p<0.05.*

We next examined whether a similar level of retention of learning was obtained across both groups. The Bootstrap test on the subtracted scores of Day 2 from Block 3 of Day 1 revealed no significant group differences (bootstrapped 95% CI [−21.370, 144.501], p=.919). An additional Bootstrap test performed on the retention test scores confirmed that both groups retained their acquired skills similarly [Mean Anaesthetised (N=15) = 53s, Mean Sham (N=16) = 46s; bootstrapped 95% CI [-11.283, 24.983], p=.419]. We also tested for the transfer of the individuation skills, and found no significant group differences [Mean Anaesthetised (N=20) = 241s, Mean Sham (N=21) = 196s; bootstrapped 95% CI [−41.971, 135.916], p=.324], suggesting that the Anaesthetised group was able to transfer the individuation skills learned on Day 1 to similar tasks’ settings as effectively as the Sham group. Together, these findings suggest that, once learned, the individuation skills were retained and transferred similarly by the two groups.

### Lower cognitive demands during motor execution

We also examined differences in skill learning by increasing cognitive task demands using a simultaneous counting task (Figure 2A). Whilst there was no difference in the proportion of incorrect answers given by each group on the counting task (bootstrapped 95% CI [−0.16688, 0.23674], p=.783), we found that the increased cognitive load uncovered a deficit in motor performance induced by the anaesthetic intervention. A direct comparison of performance on the Dual Task was significant [Mean Anaesthetised (N=19) = 71s, Mean Sham (N=19) = 41s; bootstrapped 95% CI [5.579, 39.263], p=.008], indicating an increased interference effect of the primary task. In other words, with attenuated sensory feedback, motor performance may reach less automaticity and require higher-order resources. It is important to note that no group differences were observed when comparing the difference scores between the Dual Task and Block 3 (bootstrapped 95% CI [−7.473, 32.211], p=.277), indicating that this observed deterioration of performance in the Anaesthetised group might have already been present, though latent, regardless of the cognitive load task.

### More effective retention and transfer of hand-robot coordination learning

A unique challenge for motor augmentation is the need to coordinate the movements of the robotic body part synchronously with our biological body. As shown in Figure 2B, the anaesthetic intervention did not significantly affect the acquisition of hand-robot coordination skills. To assess the acquisition of coordination skills across groups, we first ran a mixed ANOVA with Training Time (all training blocks of Day 1: Blocks 1, 2 and 3) and Group (Anaesthetised and Sham) as predictors of performance. Here, we found a statistically significant main effect of the Training Time, F(2,82)=186.242, p<.001, confirming improvements in performance over the training day, as expected. However, we did not observe a significant main effect of Group (F(1,41)=.231, p=.634), or a significant interaction (F(2,82)=.934, p=.380), suggesting that the two groups showed similar performance during the first stages of motor learning, independently of the anaesthetic intervention.

*We* did find, however, that the anaesthetic intervention impacted the retention of the acquired coordination skills. For this purpose, we ran a mixed ANOVA with Day (the final training block of Day 1 and the testing block on Day 2) and Group (Anaesthetised and Sham) as the independent variables. The ANOVA revealed a significant interaction (F(1,42)=15.429, p<.001), and no significant main effects of Day (F(1,42)=2.269, p=.139), or Group (F(1,42)=1.210, p=.278). Confirmatory paired samples t-tests indicated that the Sham group improved from Day 1 to Day 2 (t(21)=4.090, p=.001), whereas the Anaesthetised group did not (t(21)=1.620, p=.120). This resulted in a significant group difference on Day 2 (t(42)=2.285, p=.027), which indicates that the Sham group retained the hand-robot coordination skills more effectively than the Anaesthetised group over the ~24 hours interval. Furthermore, the anaesthetic intervention impaired the transfer of coordination skills, resulting in a worse performance in the Anaesthetised group on the Transfer Coordination task relative to the Sham group (t(41)=-2.10, p=.042).

### No benefits to acquisition and learning of hand-robot collaboration skills

Finally, to complete the picture of hand-robot motor interactions, we examined the impact of the anaesthetic intervention on the ability to use the Third Thumb in close collaboration with the biological fingers to grip, lift and transport objects, while maintaining constant pressure. This task contains elements of coordination, but in addition also allows for increased reliance on sensorimotor control of the biological fingers, that maintain constant pressure on the object to afford the joint grip with the Thumb. Under such cooperating conditions, we did not find any significant impact of the anaesthetic intervention on task performance and learning.

To assess the acquisition of collaboration skills, we first ran a mixed ANOVA with Training Time (all training blocks of Day 1: Blocks 1, 2 and 3) and Group (Anaesthetised and Sham groups) as predictors of performance. Here, we found a significant main effect of Training Time (F(2,78)=114.749, p<.001), but no significant main effect for Group (F(1,39)=.044, p=.834), or an interaction (F(2,78)=.116, p=.884), suggesting the groups did not improve differently across the training day. To test for group differences in retention, we ran a mixed ANOVA with Day (the final training block of Day 1 and the testing block on Day 2) and Group (Anaesthetised and Sham) as the independent variables. We found a significant main effect of Day (F(1,29) = 6.717, p=.015), but no significant effects for Group (F(1,29)=.003, p=.960) or for the interaction (F(1,29)=.320, p=.576). Overall, these results suggest that the skills needed to perform the Collaboration task were acquired and retained similarly by the two groups.

### Different toe motor control despite similar task performance during hand-robot Collaboration

It is possible that while overall performance appears similar between the two groups, the underlying strategies for achieving the same skill level are divergent. We therefore investigated differences in how the two groups used the pressure sensors to control the Third Thumb at the very first stages of motor learning. We focused on performance during Block 1 of the Collaboration task, where, as seen in Figure 2C, no performance differences were apparent. We found that the motor action patterns shown by the Anaesthetised group to achieve the same scores as the Sham group were less lateralised.

Due to violations of the assumptions of normality, group differences were analysed with bootstrap tests. Both groups did not differ in the amount of average force applied during flexion/extension and abduction/adduction movements (Flexion: Mean Sham (N=17) = 0.691 ± 0.10, Mean Anaesthetised = 0.710 ± 0.09, bootstrapped 95% CI: [−0.045, 0.081], p = .555; Abduction: Mean Sham (N=17) = 0.525 ± 0.16, Mean Anaesthetised (N=17) = 0.544 ± 0.12, bootstrapped 95% CI: [−0.074, 0.111], p = .685). However, the Anaesthetised group spent a significantly longer proportion of time making bilateral movements (movements including the simultaneous flexion and abduction of the Third Thumb) – Mean Sham (N=16) = 14.5% ± 14.3%, Mean Anaesthetised (N=20) = 27.8% ± 24%, bootstrapped 95% CI: [0.011, 0.258], p=.033. Further analyses show that this can be attributed to the Anaesthetised group spending a larger proportion of the task time making adduction movements (Mean (N=20) = 35.5% ± 33.9%), when compared to the Sham group (Mean (N=17) = 18.7% ± 18.6%, bootstrapped 95% CI: [0.006, 0.342], p=.043). The proportion of the task time spent making flexion movements did not differ between the Anaesthetised (Mean (N=19) = 48.6% ± 11.1%; and Sham (Mean (N=17) = 50.4% ± 12.6%) groups – bootstrapped 95% CI: [−0.096, 0.055], p=.636). These findings indicate that even when a task can be equally performed with attenuated sensory feedback from the Thumb controllers, this is achieved through arguably more complex and energy-consuming control patterns.

## DISCUSSION

We tested if pressure feedback from the body part proportionally controlling an extra robotic finger (Third Thumb) can support motor learning, by being used as a proxy for inferring the device state and position (device proprioception). Using local anaesthesia in a placebo-controlled design, we show that attenuating somatosensory inputs from the device controllers results in impoverished motor control and learning. Deficits in motor performance were found both during skill acquisition across training, and at the delayed retention and transfer tests. Importantly, our findings also demonstrate that despite these motor learning deficits, participants were able to learn to control the Thumb even when these pressure signals were not available. Most strikingly, performance was not impaired by anaesthesia when tasks involved close collaboration with the biological fingers, indicating that the brain could ‘close the gap’ of the missing device state and position cues by alternative means, including through continuous task-relevant somatosensory feedback from other body parts involved in the task. Nevertheless, the impairments in multitasking and the somewhat less economical control patterns underlying the performance of the Anaesthetised group highlight the unique advantages that intrinsic pressure feedback from the device controller can provide for more automatic and efficient control. Together, our findings indicate that there are multiple available avenues – involving somatosensory signals from both the controller and collaborating body-parts – that could be harnessed to increase the bidirectionality of artificial limbs control.

Initial deficits in skill acquisition were seen in the Individuation task, which required dexterous control of the Third Thumb in isolation from the biological fingers, highlighting the benefits of supplementary sensory feedback when extra precision is needed (Blank et al., 2008). Such early deficits, however, resolved by the end of the training sessions. Moreover, this effect was not observed in the Collaboration and Coordination tasks, which both required using the Thumb in conjunction with the other fingers, likely allowing for an increased reliance on sensorimotor control of the biological hand for task success (Zhu et al., 2019). These findings suggest that, during the initial stages of motor learning, other sensory cues (e.g. visual and auditory inputs from the Thumb, as well as somatosensory inputs from the collaborating hand) can probably sufficiently compensate for the lack of pressure feedback from the controllers of the device.

Nevertheless, such increased reliance on complementary feedback modalities appears to cost more pronounced cognitive resources for motor control, at least during the early stages of learning. This is well evidenced by the impairments shown by the Anaesthetised group on a Dual Task, requiring participants to multi-task by simultaneously carrying out the Individuation task and numerical operations (Witteveen et al., 2012; Guthrie et al., 2019). When fewer executive resources were available due to the need to prioritise the numerical task, the poorer motor performance in the Anaesthetised group resurfaced. This finding suggests that automaticity in device control can more readily be achieved when users can rely on task-intrinsic pressure feedback. A related finding on the distinct control patterns characterising the performance of the two groups was revealed in the Collaboration task. Here, we uncovered a potentially redundant, and thus less efficient, use of the pressure sensors to operate the Third Thumb by the Anaesthetised users in the early stages of learning. Specifically, to achieve equal task performance as the Sham group, Anaesthetised participants used more complex and energy-consuming bilateral movements. This finding shows that even when the advantages afforded by the controllers’ pressure feedback are not reflected in task performance, they provide opportunities for optimisation that are not as readily available to users relying only on other sensory cues.

Further clues for the potential advantages of surrogate state and position cues for motor learning were revealed during retention and transfer. Performance on the retention tests, which involved the repetition of the practiced tasks on the following day, reflects the relative strength of the motor memory representation over time, whereas performance on the transfer tests reflects its flexibility. Here, the impact of practice with reduced pressure feedback was evident for the Coordination task, where the Sham group continued to show improvements whereas the Anaesthetised group did not, even if toes sensitivity was by then restored for both groups. It is possible that the pressure feedback from the Thumb controllers during task training may have enabled a more accurate internal model of the motor plan, thereby contributing to a more effective error-based learning (Wolpert & Ghahramani, 1995). This may, in turn, have led to a more effective acquisition of the task, resulting in a more robust and flexible motor memory representation. Consistent with that, we have previously shown that trained participants can complete the coordination task even with no visual feedback (Kieliba et al., 2021), suggesting that users can develop a sense of position of the Thumb relative to the fingers relying mostly on somatosensory information. No group differences in retention or transfer were observed for the Individuation and Collaboration tasks, highlighting the fact that the relative role of intrinsic pressure feedback for motor learning likely depends on the usability of complementary sensory cues in the different tasks (e.g. vision and/or continuous sensorimotor feedback from the collaborating fingers). It is important to consider that both groups were receiving normal pressure feedback during the retention and transfer tests of Day 2. Therefore, it is possible that although participants learnt effectively to perform the Coordination task without pressure feedback, the incoming somatosensory inputs on Day 2 may have required them to re-learn how to integrate this information into their internal model. It was impossible for us to empirically test the theory that under anaesthesia retention would have been more complete, due to unsafe toxicity build-up that could result from consecutive-day anaesthetic blocks. Nevertheless, considering that performance during Day 2 did not vary across groups for the other tasks, we believe this interpretation is less likely.

Our findings bear important implications for the development of assistive and augmentative motor interfaces. It is well recognised that some of the remaining functional inadequacy of BMIs derives from the lack of somatosensory feedback, including both touch and proprioception. Akin to deafferented patients and the anaesthetised participants in our study, BMI-controlled movements require considerable attention and do not typically achieve near-natural levels of dexterity and fluidity (Tomlinson & Miller, 2016). Here, we show that leveraging task-intrinsic somatosensory inputs can substantially enhance motor performance and learning with artificial robotic limbs. By reading motor commands from the cortex and bypassing completely the body, current BMIs may be missing important opportunities for harnessing such task-intrinsic somatosensory signals for motor control. Moreover, engaging the body in some form of device control may not only optimise motor control through increasing task-relevant sensory feedback, but also serve rehabilitation by providing physical activity, with the potential to prevent muscle atrophy and maintain any residual mobility (Pierella et al., 2015). In light of the advantages documented here, we suggest that BMI systems would probably benefit from exploring a hybrid approach, where neuronal recordings are coupled with some form of bodily engagement. BMI systems that restore movement through neuromuscular stimulation of the patient’s own limb (Bouton et al., 2016, Ajiboye et al., 2017; Bockbrader et al., 2019; Ganzer et al., 2020) may already be benefiting from such opportunities.

## ACKNOWLEDGEMENTS

This work was supported by an ERC Starting Grant (715022 EmbodiedTech) and by Sir Halley Stewart Charitable Trust (580), awarded to TRM, who was further funded by a Wellcome Trust Senior Research Fellowship (215575/Z/19/Z).

